# Genetic dependency of atrial fibrillation-associated risk genes across tissue types: Discovering novel therapeutic targets

**DOI:** 10.64898/2026.06.11.731776

**Authors:** Vishal M. Bommineni, Ubaldo G. Morales, Ziran Yang, Zachery Lerch, Monicka Felix, Rias Ali

## Abstract

**Background:** Addressing the underlying causes of atrial fibrillation (AFib) is critically important. While potential AFib-related genes have been recognized, the impact of modifying these genes in humans remains poorly understood.

**Objective:** We assessed the cellular dependencies of 309 genes previously associated with AFib through genome-wide association studies using data from the Cancer Dependency Map project, aiming to prioritize potential therapeutic targets with minimal off-target effects.

**Methods:** We analyzed CRISPR-Cas9 knockout (CHRONOS scores) and RNA interference (RNAi) knockdown (DEMETER2 scores) screening data from 1,927 human cell lines across 24 tissue types, focusing on tissues associated with AFib initiation, presentation, and progression: autonomic ganglia, central nervous system (CNS), and soft tissue. We examined the expression and dependency scores of the AFib-associated genes, identifying significant correlations between gene expression and cellular dependency within specific tissues using Pearson correlation coefficients and controlling the false discovery rate (FDR) at 5%.

**Results:** Out of the 309 AFib-associated genes, 206 genes (66.7%) had CHRONOS dependency scores and 229 (74.1%) had DEMETER2 dependency scores available. Several genes showed significant negative dependency scores (CHRONOS < −0.5) across multiple tissues, indicating potential off-target effects if inhibited. In contrast, we identified 12 genes with significant expression-driven dependencies within AFib-associated tissues. In CNS cell lines, *HAND2* (R = −0.456, FDR = 0.002) and *VGLL2* (R = −0.434, FDR = 0.005) showed significant negative correlations between gene expression and cellular dependency. In soft tissue cell lines, *BEST3* (R = −0.679, FDR = 0.001) and *PITX2* (R = −0.679, FDR = 0.001) also demonstrated strong negative correlations. Additionally, *ERBB4* in CNS lines showed a significant negative correlation (R = −0.361, FDR = 0.048). These findings suggest that inhibiting these genes may selectively affect high-expressing cells in AFib-associated tissues while minimizing effects on other tissues.

**Conclusion:** Our analysis identified *HAND2*, *VGLL2*, *BEST3*, and *ERBB4* as potential therapeutic targets for AFib, demonstrating significant expression-driven dependencies in AFib-associated tissues with no pan-tissue essentiality. These results provide a quantitative basis for developing targeted therapies with reduced off-target effects.

**CONDENSED ABSTRACT:** Atrial fibrillation (AFIB) is one of the most common cardiac arrhythmias with numerous known risk factors. Although many AFIB-associated genes have been identified, the impact of screening or the effects of modifying these genes in humans remain poorly understood. We examined CRISPR knockout and RNAi knockdown screen data from nearly 2,000 human cell lines to assess the cellular dependencies of 309 genes associated with AFIB, previously identified through genome-wide association studies. Some genes demonstrate broad cell dependencies across various tissue types, indicating potential off-target effects if inhibited. Conversely, *HAND2*, *VGLL2*, *BEST3*, and *ERBB4* were identified as genes of interest because their genetic knockouts specifically impacted high-expressing cells from tissue lineages pertinent to AFIB and/or were not pan-dependent. Overall, analyses of genetic screen data identified AFIB-associated genes whose knockout or knockdown selectively affected cell lines of relevant tissue lineages, prioritizing targets for potential AFIB treatments.

## INTRODUCTION

Atrial fibrillation (AFIB) is the most common sustained cardiac arrhythmia worldwide, affecting more than 60 million individuals and substantially increasing the risk of stroke, heart failure, and mortality^1,2^. Despite major advances in rhythm control and ablation therapies, AFIB management remains challenging due to its multifactorial etiology and incomplete understanding of the underlying molecular mechanisms.

Genetic predisposition plays a central role in AFIB susceptibility. Large-scale genome-wide association studies (GWAS) have identified more than 100 loci linked to AFIB, including well-established genes such as *PITX2*, *ZFHX3*, *KCNN3*, *PRRX1*, *TBX5*, and *KCNH2*, which are involved in cardiac development, ion channel regulation, and atrial structural remodeling^3,4,10,16–20,23,24,47^. However, these variants explain only a modest fraction of AFIB heritability, and the functional consequences of most risk genes remain unclear^7,8,21^.

Early-onset AFIB, particularly in patients under 45 years of age and without conventional risk factors, is often enriched for monogenic or highly penetrant variants, sometimes overlapping with genes implicated in channelopathies and cardiomyopathies such as *SCN5A*, *LMNA*, and *TTN*^27,36,47.^ Yet, most cases of adult-onset AFIB arise from complex polygenic backgrounds interacting with structural and environmental contributors such as hypertension and atrial dilation. The fact that not all individuals with similar clinical risk factors develop AFIB underscores the contribution of genetic modifiers that remain to be functionally characterized^8,21,47^.

While GWAS has identified loci associated with AFIB, these studies primarily capture statistical associations rather than causal mechanisms. Accordingly, functional genomic analyses can serve as prioritization tools to refine candidate genes for further study rather than to establish definitive causal mechanisms. Understanding how these genes influence cellular physiology, particularly whether they are essential for cell survival, signaling, or excitability in relevant tissues, may help prioritize candidates for deeper mechanistic investigation and therapeutic exploration.^5,6,13,14^.

Recent advances in functional genomics, including genome-wide CRISPR-Cas9 knockout and RNA interference (RNAi) knockdown screens, enable systematic assessment of gene essentiality across hundreds of human cell types^5,6,29,30,33^. The Cancer Dependency Map (DepMap) aggregates such data from nearly 2,000 human cell lines, providing a powerful resource to evaluate gene function and dependency in diverse biological contexts (https://depmap.org/portal) ^6,29^. Although DepMap was designed for oncology research, its large-scale dataset allows exploration of genetic dependencies in tissue lineages relevant to other diseases, including cardiac and neural cells involved in AFIB pathogenesis^6,47^.

In this study, we analyzed 309 AFIB-associated genes that were identified through prior GWAS^3,4,7,10,16–20,23,26^ and evaluated their cellular dependencies using DepMap’s genome-wide CRISPR (CHRONOS) and RNAi (DEMETER2) datasets^29,30,33^. By integrating gene expression and dependency data across 24 tissue lineages, including autonomic ganglia, central nervous system, and soft tissue, we aimed to prioritize AFIB risk genes whose inhibition selectively affects relevant cell types while minimizing off-target effects. This approach provides a functional layer atop association data, helping to prioritize AFIB-linked genes that may represent viable, tissue-selective therapeutic targets.

## METHODS

An overview of the methods are visualized in Fig. 1.

**Figure 1.**
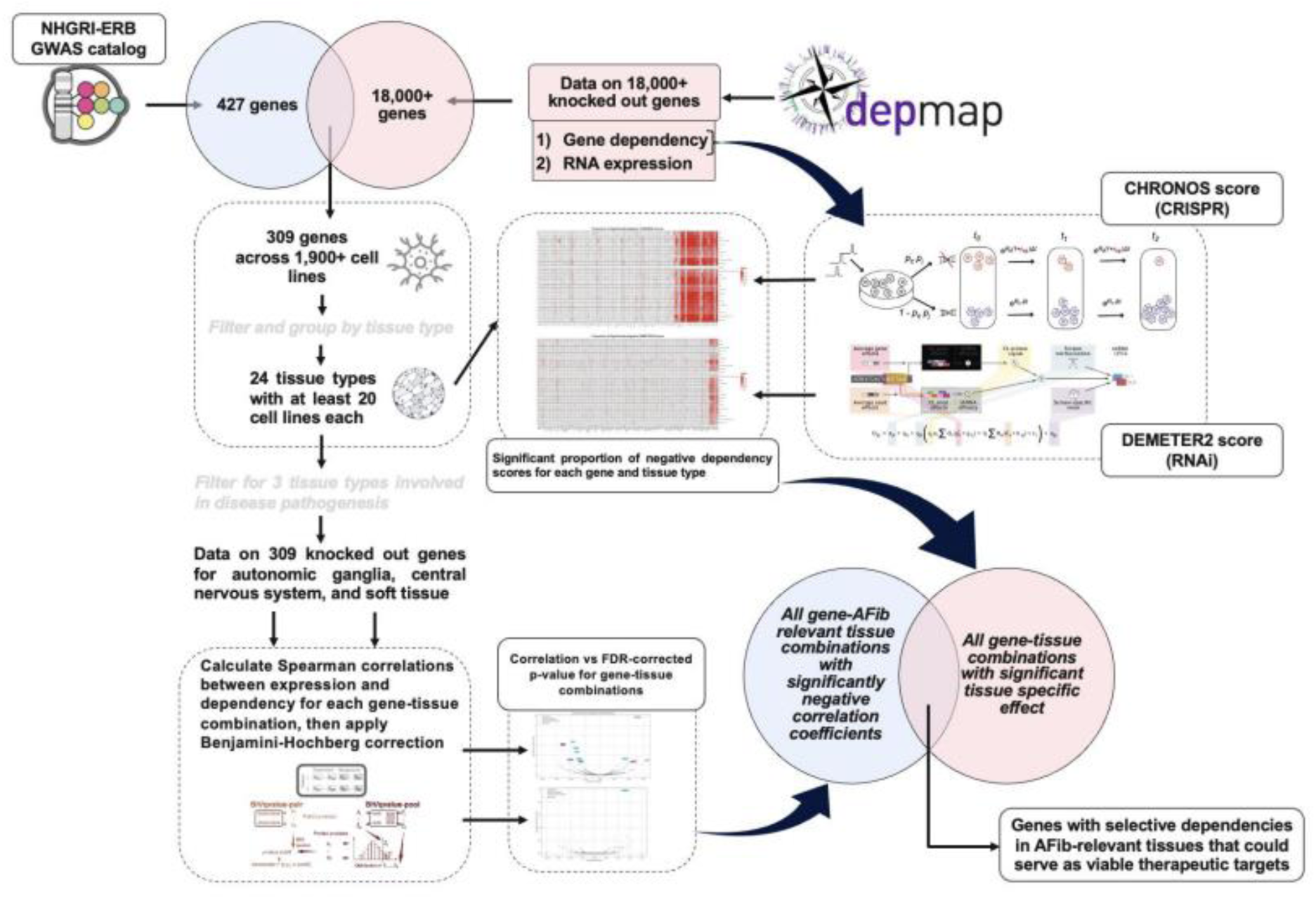
Overview of methodology. All computational analyses were performed in Python (version 3.11.0).

### Data Source and Download

Genes used in this study were obtained from the NHGRI-ERB GWAS catalog, a curated collection of all human genome-wide association studies published in peer-review journals^7^. 427 unique genes from GWAS^3,4,8–28^ were compiled. For each gene, cancer gene expression and dependency scores were downloaded from the DepMap Portal Public 24Q2 release in July 2024^29^; at least one of these data were available for 309 genes.

### Expression-driven dependency using DepMap

The DepMap Public 24Q2 release contains the results of genome-wide CRISPR knockout screens for 18,436 genes across 1927 cell lines, including both normal and cancer cell lines, as well as the RNA expression of 19,098 genes across 1927 cell lines. 24 tissue types from DepMap with at least 20 cell lines each were selected for further analysis from an original list of 66 tissue types. For each of the 309 genes associated with AFIB, those with both gene expressions and corresponding CHRONOS dependency scores were analyzed. CHRONOS is an open-source algorithm that uses a cell population dynamics model to infer gene knockout fitness effects in CRISPR screens^30^; the CHRONOS score reflects the change in cell proliferation upon the CRISPR knockout of the respective gene in a particular cell line. For each AFIB-associated gene, we calculated the proportion of significantly negative CHRONOS scores (< −0.5) by tissue type to generate heatmaps. A negative CHRONOS score indicates that a gene knockout results in a slower growth rate of a cell line, with a score less than −0.5 indicating a notable reduction^31^, although it is not deterministic.

We also calculated the Pearson correlation and corresponding p-value for each AFIB-associated gene after stratifying the gene expression and CHRONOS scores by tissue type. This analysis was limited to examining expression-driven dependencies in three tissue types: autonomic ganglia, central nervous system, and soft tissue for their involvement in AFIB initiation, presentation, and progression. Using the Benjamini & Hochberg procedure^32^ to control the false discovery rate (FDR) of a family of hypothesis tests, Spearman correlation coefficients were calculated for each gene-tissue combination. Volcano plots were plotted with the correlation coefficients (x-axis) against the negative log of FDR-corrected p-values (y-axis). We then focused on gene-tissue combinations with negative correlation coefficients as a high expression of an AFIB-associated gene and accompanying low CHRONOS score indicates that the gene is needed for cancer cell survival in knockout experiments. Scatterplots of dependency versus expression were also plotted for AFIB gene-tissue combinations with the highest negative log p-values.

The DepMap release also contains genetic dependency score estimates from RNAi loss-of-function screens for 16,837 genes across 1927 cell lines. These scores were calculated using DEMETER2^33^, a framework developed by McFarland et al. to unify results from multiple large-scale RNAi screening datasets. Calculations of correlations, volcano plots highlighting those correlations, and scatterplots of AFIB-associated genes were completed in a similar process to that of the CHRONOS scores. Analysis of genetic dependency data was performed using Python 3.11.0.

## RESULTS

### Gene list and DepMap data assembly

A curated list of 309 AFIB-associated genes was considered for this study (Supplementary Table S1). Of these 309 genes, 304 genes had expression data, with 206 of those genes (67.8%) having at least one non-zero CHRONOS dependency score and 229 of those genes (75.3%) having at least one non-zero DEMETER2 dependency score available across the included tissue types. There were 24 tissue types with data for at least 20 cell lines each from the DepMap 2024 Q2 release which were selected for further analysis (Table 1). We specifically highlighted results from tissue types most relevant to AFIB etiology: autonomic ganglia (AG), central nervous system (CNS), and soft tissue (ST).

**Table 1.** Overview of tissue types and the number of unique cell lines used for cell line genetic dependency analysis.

| Tissue Name | Number of Cell Lines |
| --- | --- |
| Pleural Effusion | 183 |
| Hematopoietic and Lymphoid Tissue | 171 |
| Lung | 149 |
| Lymph Node | 135 |
| Central Nervous System | 132 |
| Ascites | 94 |
| Skin | 85 |
| Upper Aerodigestive Tract | 75 |
| Soft Tissue | 70 |
| Bone | 66 |
| Bone Marrow | 66 |
| Liver | 56 |
| Kidney | 56 |
| Breast | 50 |
| Ovary | 41 |
| Large Intestine | 40 |
| Biliary Tract | 38 |
| Fibroblast | 36 |
| Endometrium | 36 |
| Urinary Tract | 34 |
| Pancreas | 34 |
| Esophagus | 34 |
| Colon | 29 |
| Autonomic Ganglia | 22 |

### Pan-tissue genetic dependency

To examine the genetic dependency of AFIB-associated genes across tissues, we identified the proportion of cell lines within each tissue type showing significantly negative dependency scores less than −0.5, a threshold used by many other authors, with lower dependency scores indicating that a gene is required for cell survival and proliferation (Fig. 2A–B). From the 309 AFIB risk genes found through GWAS, *YWHAE, CDK6, SLC25A26, SLC9B1, SNRNP27, XPO1, EXT1, ZPR1, CDC27, HMGCR, NACA, GOSR2, SUGP1, NRBP1, PMVK, PTK2, CELSR2, POLR1F, WDR1, HMGA1, PFDN1, POLD1, UBE2D3, E4F1,* and *PSMB7* had varying levels of negative CHRONOS scores ranging from 1% to 87% across nearly all tissue types. *ARL5B, ASAH1, BCL3, CFL2, FAM76B, FGF5, IL6R, KCNH2, LHX3, MICU2, PRRX1, REEP3, RFLNA, SIK3, SOWAHD, SYNE2, TMT1B, USP3, WIPF1, ZFHX3,* and *ZHX3* had negative CHRONOS scores predominantly localized to one or more hematopoietic and lymphoid (HL), lung (LNG), pleural effusion (PE), and/or lymph node (LN) tissues. We also identified the proportion of cell lines with significant negative genetic dependency scores (DEMETER2 score < −0.5) utilizing data from large-scale RNAi screens. *XPO1, CDC27, NACA, PRDM6, WDR1, POLD1,* and *PSMB7* had varying levels of negative DEMETER2 scores from 1% to 76% across nearly all tissue types. *ADRB1*, *AKAP6*, *AOPEP, APOLD1, ATXN1, CNTN5, ESR2, FADS2, KLHL38, LIN54, METRN, PAPPA, RPL3L, SEPTIN6, SH3PXD2A, SIRT1, SMAD7,* and *TAB2* had negative DEMETER2 scores localized to one of more HL, LNG, PE, and/or LN tissues.

**Figure 2.**
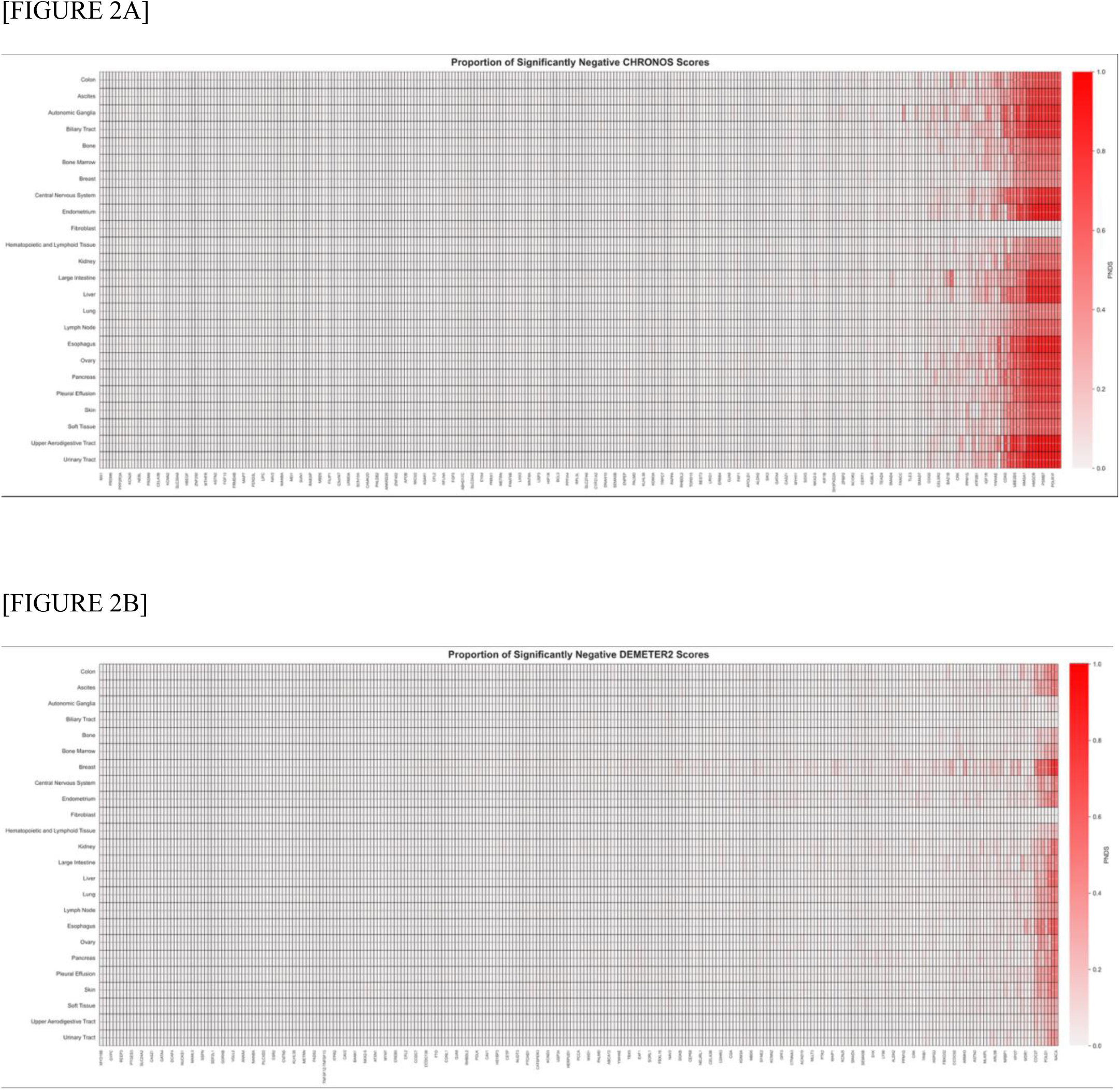
Genetic dependency of atrial fibrillation risk genes in CRISPR knockout screens across 24 tissue lineages. A) The proportion of negative dependency scores less than −0.5 in cell lines stratified by tissue types based on the CHRONOS scores derived from CRISPR knockout screens. 24 tissue types from the DepMap project with data for at least 20 cell lines were utilized to evaluate 304 atrial fibrillation risk genes (98 genes visualized based on selective- and pan-knockout determination) as potential targets. B) The proportion of negative dependency scores less than - 0.5 in cell lines stratified by tissue types based on the DEMETER2 scores derived from CRISPR knockout screens. 24 tissue types from the DepMap project with data for at least 20 cell lines were utilized to evaluate 284 atrial fibrillation risk genes (119 genes visualized based on selective- and pan-knockout determination) as potential targets. Abbreviation: PNDS = “Proportion of Negative Dependency Scores (Less Than −0.5)”.

The mostly concordant CHRONOS/RNAi screen results suggest that knocking down/out of several genes (i.e., *XPO1* and *NACA*) results in widespread consequences affecting the survival of many/most cells, whereas other candidates (i.e., *HAND2*) more selectively affected AFIB-relevant cell types, thereby potentially serving as better targets.

### Expression-driven cellular dependencies of AFIB risk genes

We reasoned that AFIB-associated genes may show aberrant expression in disease-driving and affected cells. Thus, candidate genes would likely serve as better targets if their knockout or knockdown most strongly influenced the cells showing aberrant expression of the targeted genes. We next sought to further filter for genes whose expression is significantly correlated with dependencies of cell lines within these tissue types. We conducted a systematic Pearson correlation analysis using CRISPR knockout-based CHRONOS scores and expression data to identify such genes of interest; we identified 12 genes whose expression was significantly associated (FDR < 0.05) with cellular dependencies (Table 2).

**Table 2.** Genes whose expressions are significantly associated with CHRONOS cellular dependency scores, grouped by tissue.

|  | Gene | Correlation | FDR-adjusted P-value |
| --- | --- | --- | --- |
| Central Nervous System | *CDKN1A* | 0.597 | 0.001*** |
|  | *CDK6* | -0.501 | 0.001*** |
|  | *HAND2* | -0.456 | 0.002** |
|  | *VGLL2* | -0.434 | 0.005** |
|  | *IGF1R* | -0.394 | 0.024* |
|  | *APOLD1* | 0.390 | 0.025* |
|  | *MLLT3* | -0.371 | 0.039* |
|  | *ERBB4* | -0.361 | 0.048* |
| Soft Tissue | *PITX2* | -0.679 | 0.001*** |
|  | *BEST3* | -0.671 | 0.001*** |
|  | *PHLDB2* | -0.555 | 0.030* |
| Autonomic Ganglia | *COG5* | 0.778 | 0.035* |

\* $p < 0.05$ ; \*\* $p < 0.01$ ; \*\*\* $p < 0.001$ (Benjamini–Hochberg FDR-adjusted)

We further conducted an analogous correlation analysis using RNAi knockdown-based DEMETER2 scores and expression data to identify similarities and differences to CHRONOS results. We found one gene with significant expression-driven dependency: *CDKN1A* in CNS cell lines (R = 0.59, FDR = 0.0037). Notably, the positive correlation observed for CDKN1A indicates that higher expression was associated with less negative dependency scores, consistent with a context-dependent growth-suppressive or protective role rather than essentiality for proliferation. Volcano plots for all gene-tissue combinations as well as correlation plots for the highest negative log p-values in both the CRISPR knockout and RNAi knockdown experiments are shown in Fig. 3A-D.

**Figure 3.**
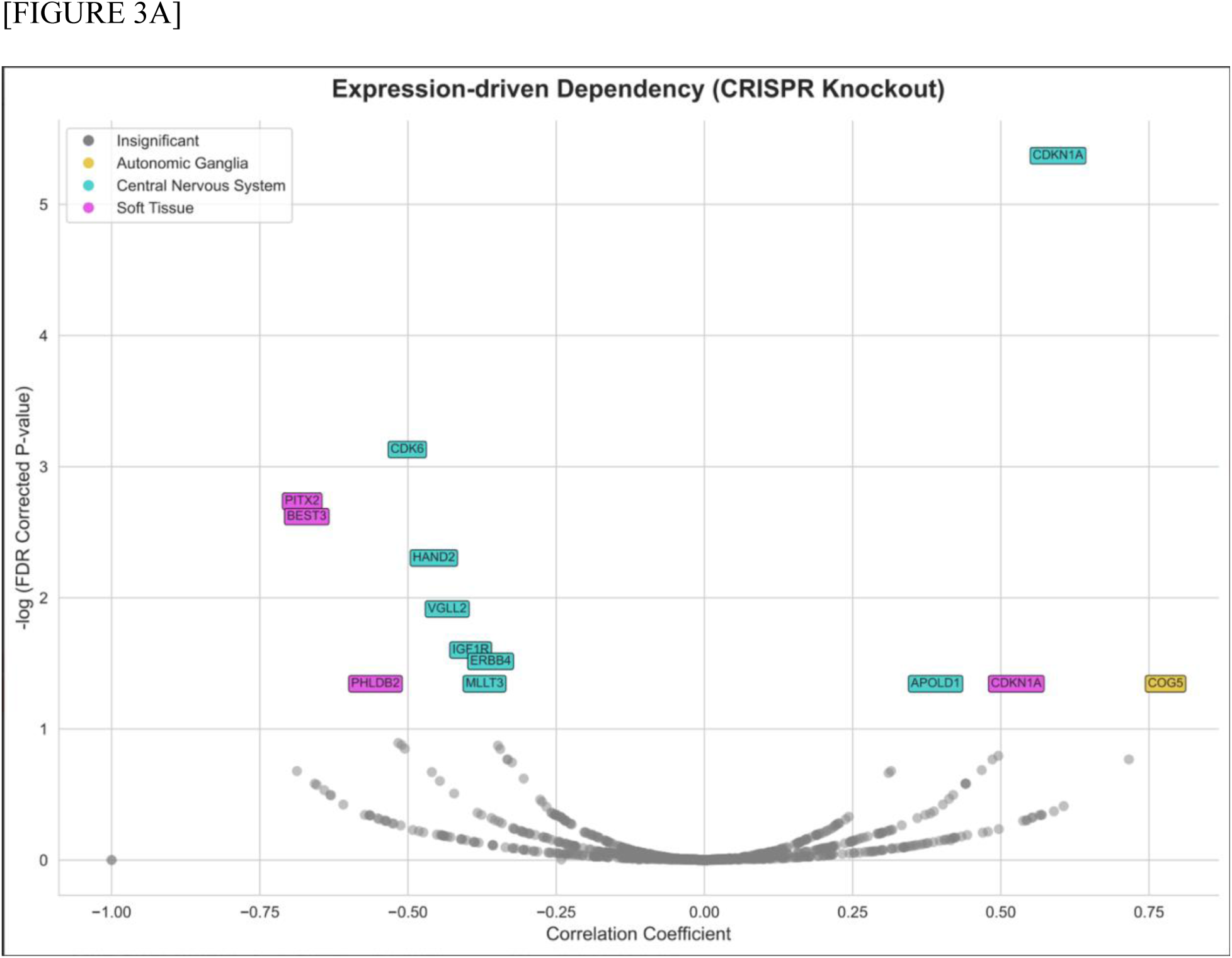
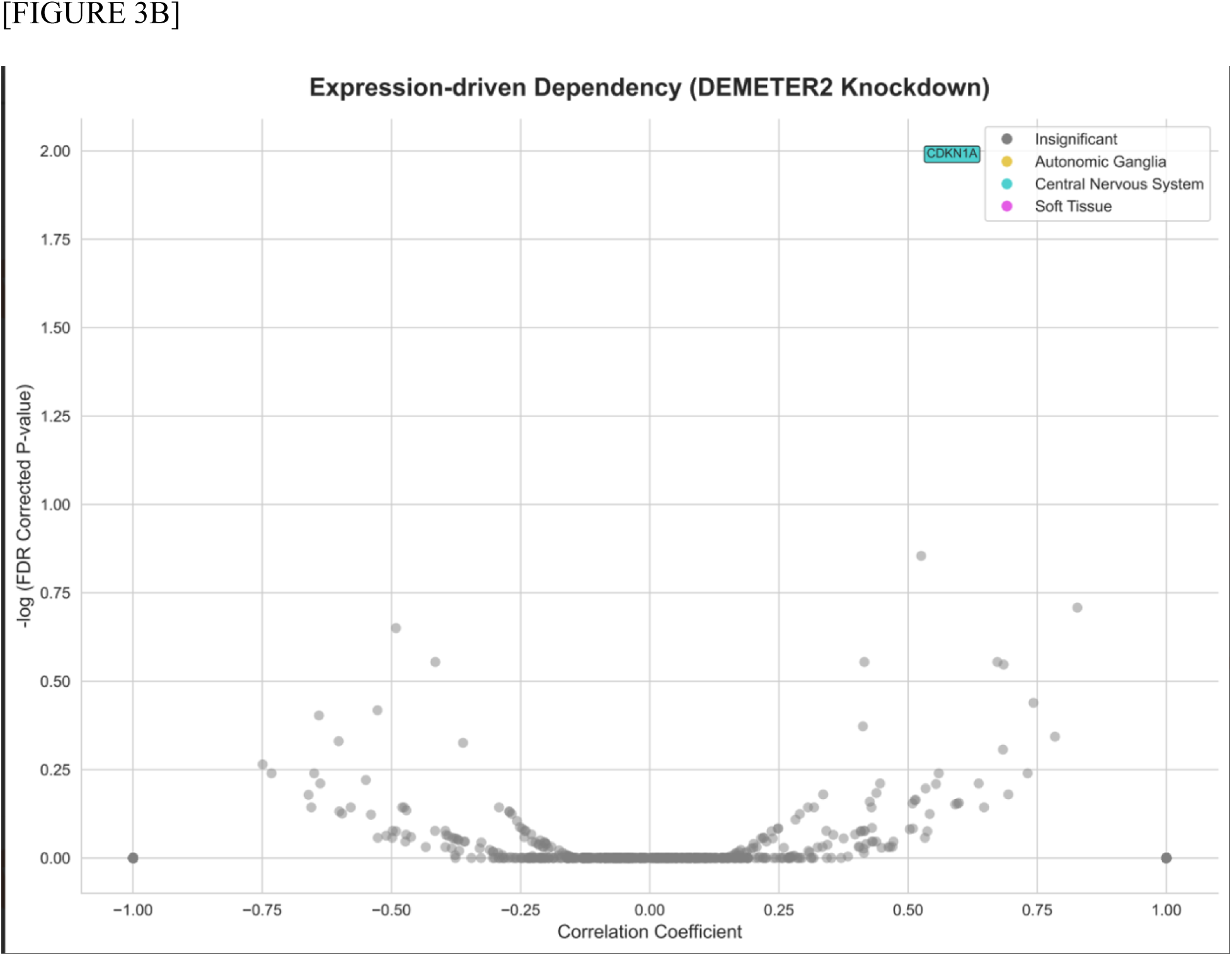
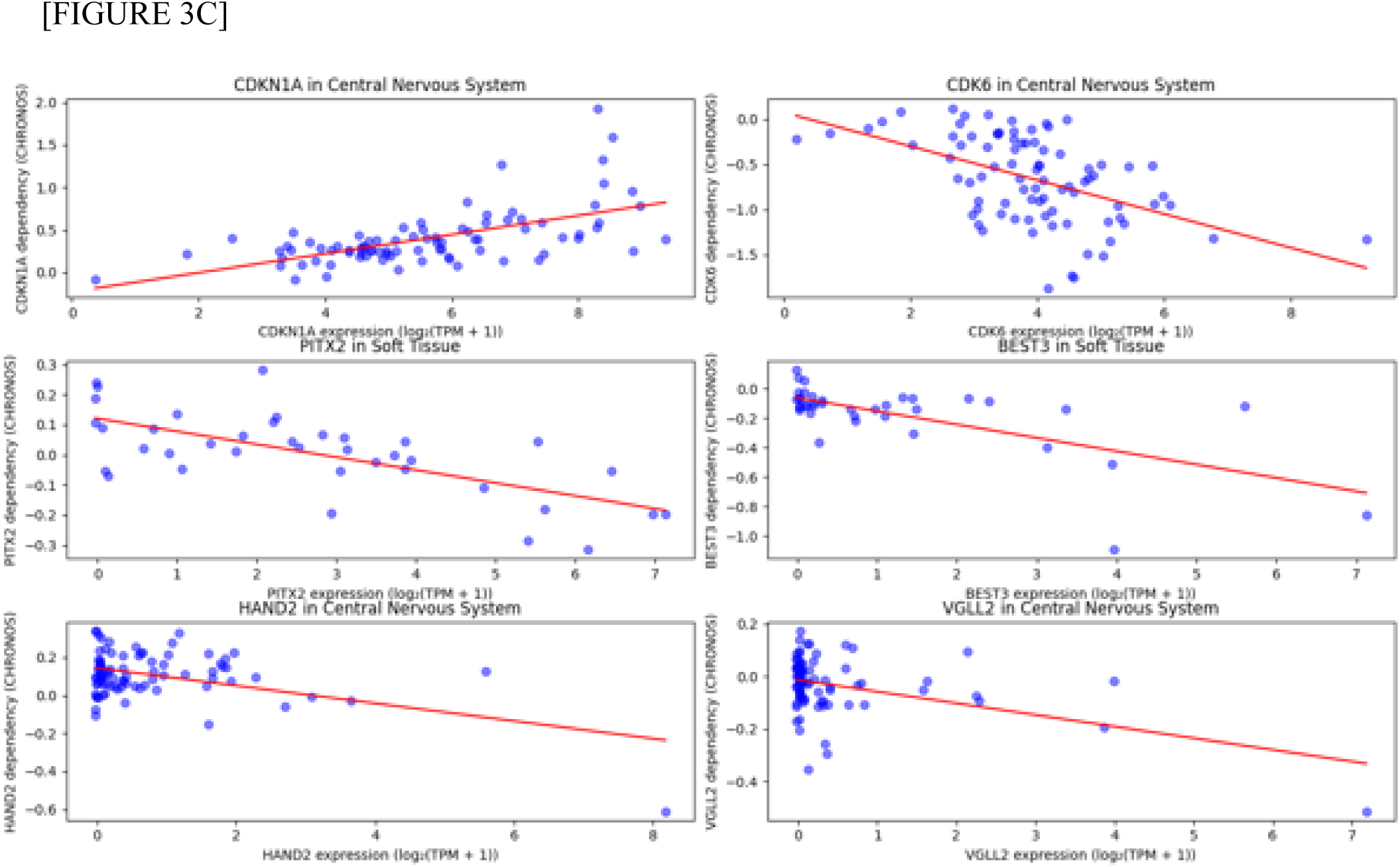
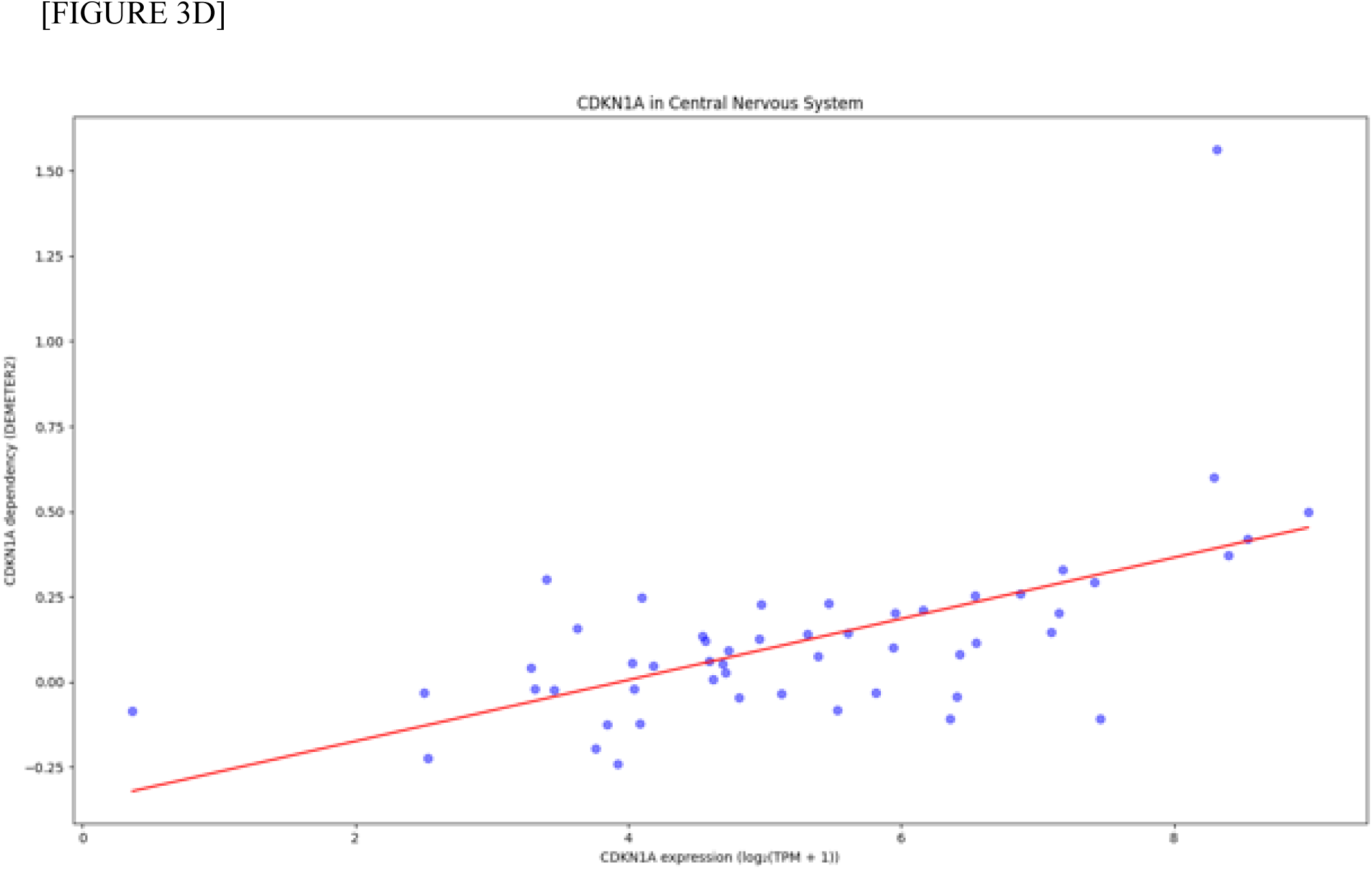
(Top to Bottom: A-D) Expression-driven dependency of atrial fibrillation (AFIB) risk genes. A) A volcano plot showing expression-driven dependency of AFIB genes based on CRISPR knockout screen data. B) A volcano plot showing expression-driven dependency of AFIB genes based on RNAi knockdown screen data. For A) and B), Pearson correlation coefficients were calculated for risk genes using a gene’s expression values versus dependency scores for four tissue types (color-coded) relevant to AFIB etiology: autonomic ganglia, central nervous system, and soft tissue. The correlation plots of the expression-driven dependency were shown for the six strongest associations found for C) *CDKN1A, CDK6, HAND2,* and *VGLL2* in CNS cell lines and *PITX2* and *BEST3* in ST cell lines against CHRONOS scores. Correlation plots of expression-driven dependency were shown for the strongest association found for D) *CDKN1A* in CNS cell lines against DEMETER2 scores. Cell lines are shown by blue points, and best-fitted regression lines are shown in red.

## DISCUSSION

This study represents, to our knowledge, the first large-scale functional genomic analysis of AFIB-associated genes across nearly 2,000 human cell lines. By integrating genome-wide CRISPR-Cas9 and RNAi dependency data with expression profiles, we sought to bridge the gap between genetic association and functional relevance, identifying genes whose inhibition affects cellular survival in AFIB-relevant tissues.

Our findings build upon prior GWAS and experimental studies that have implicated key regulators such as *PITX2*, *ZFHX3*, *KCNN3*, *PRRX1*, *HAND2*, and *TBX5* in AFIB susceptibility^3,4,10,16–20,23,24,45–47^. Many of these genes play roles in cardiac development, electrophysiologic signaling, and structural remodeling, yet their therapeutic tractability remains uncertain. Among the 309 genes examined, several of these well-established AFIB loci were included in our dataset and exhibited distinct dependency patterns. For instance, *PITX2* and *HAND2*—both central to left atrial and pulmonary venous myocardial development—showed tissue-selective dependencies, supporting their functional relevance and potential as therapeutic targets^44–46^. Conversely, widely expressed genes such as *CDK6*, *XPO1*, and *NACA* demonstrated pan-essentiality, suggesting that systemic inhibition could be deleterious^5,6,29^.

Our expression–dependency correlation analyses further highlighted *HAND2*, *VGLL2*, *BEST3*, and *ERBB4* as candidates whose knockdown selectively affected high-expressing cell lines from AFIB-relevant tissues. Notably, *HAND2* and *VGLL2* are involved in cardiac and neural crest development^43–46^, while *BEST3*, a calcium-activated chloride channel, has been linked to inflammation and endothelial signaling—processes increasingly recognized in AFIB pathogenesis^34,35,37–39^. These findings complement prior reports linking *HAND2* dysregulation to atrial structural remodeling^44–46^ and BEST3 to anti-inflammatory pathways in endothelial cells, aligning with evidence that inflammation and fibrosis contribute to AFIB initiation and persistence^34,37–39^.

Previous studies examining AFIB genetics have largely focused on association mapping and pathway enrichment, rather than functional essentiality. To date, no large-scale study has systematically screened AFIB-associated genes for cellular dependency or tissue-selective knockout sensitivity. Thus, our analysis extends the field by leveraging existing functional genomics resources to prioritize AFIB risk genes based on dependency and expression context—an approach more commonly used in oncology than cardiology^5,6,29,30,33,48^.

Importantly, our findings provide a framework for distinguishing pan-essential from selectively essential genes, which has major implications for therapeutic design. For example, genes whose knockout is lethal across most cell types (e.g., *CDK6*, *XPO1*) may pose toxicity risks if targeted systemically, whereas selectively essential genes (e.g., *HAND2*, *VGLL2*, *BEST3*) could represent safer, tissue-restricted therapeutic targets^6,29,43–46^.

Although this study uses data from primarily cancer-derived cell lines, our approach offers a filtering strategy to identify AFIB-relevant targets for further validation in cardiac-specific models, such as patient-derived induced pluripotent stem cells (iPSC)-derived atrial cardiomyocytes, cardiac organoids, or animal models. Future studies integrating single-cell transcriptomic and CRISPR screening data from human cardiac tissues could refine these predictions and help define gene networks central to atrial remodeling and arrhythmogenesis^47,48^.

From a translational perspective, prioritized genes identified in this analysis could be investigated using targeted loss- or gain-of-function approaches in human iPSC-derived atrial cardiomyocytes to assess effects on action potential duration, calcium handling, and arrhythmogenic susceptibility. Small-molecule inhibitors, antisense oligonucleotides, CRISPR-based gene modulation systems, or RNA interference platforms could be evaluated for selectively essential genes to determine whether modulation alters electrophysiologic phenotypes without inducing widespread cytotoxicity. In parallel, tissue-restricted delivery strategies, such as adeno-associated viral vectors with cardiac-specific promoters, may help mitigate off-target effects suggested by pan-essential dependency patterns. Such studies would clarify therapeutic tractability and inform the development of precision-targeted AFIB interventions.

In summary, this work integrates association genetics with functional genomics to prioritize AFIB-linked genes based on tissue-selective dependency, bridging a major gap between GWAS discovery and translational application. By identifying a subset of AFIB genes whose inhibition selectively impacts relevant cell types, this study lays the foundation for future mechanistic and therapeutic investigations targeting AFIB at its molecular roots.

## CONCLUSION

In this study, we systematically evaluated the cellular dependencies of 309 atrial fibrillation (AFIB)-associated genes by integrating genome-wide CRISPR-Cas9 and RNAi screening data from nearly 2,000 human cell lines, alongside matched gene expression data from the DepMap project^6,29–31^. Our results demonstrate that while several AFIB-associated genes—such as *XPO1*, *CDK6*, and *NACA*—exhibit pan-essentiality across diverse tissue types, and thus may present challenges as therapeutic targets due to potential systemic toxicity, other genes display more tissue-selective dependency patterns.

Specifically, genes including *HAND2*, *PITX2*, *BEST3*, *VGLL2*, and *ERBB4* showed expression-driven dependencies in cell types relevant to AFIB pathogenesis, namely the central nervous system, autonomic ganglia, and soft tissue^20,27,35,36,40–43^. These findings suggest that targeting such genes may enable tissue-selective therapeutic strategies, with potentially fewer off-target effects compared to pan-essential genes.

Although a limitation of this study is the reliance on cell lines derived primarily from cancer tissues—which may not fully reflect AFIB-relevant cardiac cell types—the use of large-scale functional genomic datasets offers a powerful filtering framework. It allows for the prioritization of genes for downstream validation in more disease-relevant systems, such as iPSCs, cardiac organoids, or in vivo AFIB models^4,32,33^. Ultimately, our findings contribute to the identification of potential therapeutic targets and improve understanding of the molecular mechanisms underlying AFIB.

## HCA HEALTHCARE DISCLAIMER

This research was supported (in whole or part) by HCA Healthcare and/or an HCA Healthcare affiliated entity. The views expressed in this presentation represent those of the author and do not necessarily represent the official views of HCA Healthcare or any of its affiliated entities.

## ACKNOWLEDGEMENTS

N/A

## ABBREVIATIONS

N/A

## REFERENCE LIST

1. Chugh, S.S., et al. Worldwide epidemiology of atrial fibrillation: a Global Burden of Disease 2010 Study. Circulation 129, 837–847 (2014).

2. Kirchhof, P., et al. 2016 ESC Guidelines for the management of atrial fibrillation developed in collaboration with EACTS. Eur Heart J 37, 2893–2962 (2016).

3. Roselli, C., et al. Multi-ethnic genome-wide association study for atrial fibrillation. Nat Genet 50, 1225–1233 (2018).

4. Nielsen, J.B., et al. Biobank-driven genomic discovery yields new insight into atrial fibrillation biology. Nat Genet 50, 1234–1239 (2018).

5. Meyers, R.M., et al. Computational correction of copy number effect improves specificity of CRISPR-Cas9 essentiality screens in cancer cells. Nat Genet 49, 1779–1784 (2017).

6. Arafeh, R., Shibue, T., Dempster, J.M., Hahn, W.C. & Vazquez, F. The present and future of the Cancer Dependency Map. Nat Rev Cancer (2024).

7. Sollis, E., et al. The NHGRI-EBI GWAS Catalog: knowledgebase and deposition resource. Nucleic Acids Res 51, D977–D985 (2023).

8. Emmert, D.B., et al. Genetic and Metabolic Determinants of Atrial Fibrillation in a General Population Sample: The CHRIS Study. Biomolecules 11(2021).

9. Carcel-Marquez, J., et al. A Polygenic Risk Score Based on a Cardioembolic Stroke Multitrait Analysis Improves a Clinical Prediction Model for This Stroke Subtype. Front Cardiovasc Med 9, 940696 (2022).

10. Gudbjartsson, D.F., et al. Variants conferring risk of atrial fibrillation on chromosome 4q25. Nature 448, 353–357 (2007).

11. Jiang, L., Zheng, Z., Fang, H. & Yang, J. A generalized linear mixed model association tool for biobank-scale data. Nat Genet 53, 1616–1621 (2021).

12. Kertai, M.D., et al. Genome-wide association study of new-onset atrial fibrillation after coronary artery bypass grafting surgery. Am Heart J 170, 580–590 e528 (2015).

13. He, L., et al. Pleiotropic Meta-Analyses of Longitudinal Studies Discover Novel Genetic Variants Associated with Age-Related Diseases. Front Genet 7, 179 (2016).

14. Walters, R.G., et al. Genotyping and population characteristics of the China Kadoorie Biobank. Cell Genom 3, 100361 (2023).

15. Hong, M., et al. Ethnic similarities in genetic polymorphisms associated with atrial fibrillation: Far East Asian vs European populations. Eur J Clin Invest 51, e13584 (2021).

16. Christophersen, I.E., et al. Large-scale analyses of common and rare variants identify 12 new loci associated with atrial fibrillation. Nat Genet 49, 946–952 (2017).

17. Gudbjartsson, D.F., et al. A sequence variant in ZFHX3 on 16q22 associates with atrial fibrillation and ischemic stroke. Nat Genet 41, 876–878 (2009).

18. Benjamin, E.J., et al. Variants in ZFHX3 are associated with atrial fibrillation in individuals of European ancestry. Nat Genet 41, 879–881 (2009).

19. Ellinor, P.T., et al. Common variants in KCNN3 are associated with lone atrial fibrillation. Nat Genet 42, 240–244 (2010).

20. Low, S.K., et al. Identification of six new genetic loci associated with atrial fibrillation in the Japanese population. Nat Genet 49, 953–958 (2017).

21. Weng, L.C., et al. Heritability of Atrial Fibrillation. Circ Cardiovasc Genet 10(2017).

22. Larson, M.G., et al. Framingham Heart Study 100K project: genome-wide associations for cardiovascular disease outcomes. BMC Med Genet 8 Suppl 1, S5 (2007).

23. Nielsen, J.B., et al. Genome-wide Study of Atrial Fibrillation Identifies Seven Risk Loci and Highlights Biological Pathways and Regulatory Elements Involved in Cardiac Development. Am J Hum Genet 102, 103–115 (2018).

24. Ellinor, P.T., et al. Meta-analysis identifies six new susceptibility loci for atrial fibrillation. Nat Genet 44, 670–675 (2012).

25. Wang, X., et al. Common and rare variants associated with cardiometabolic traits across 98,622 whole-genome sequences in the All of Us research program. J Hum Genet 68, 565–570 (2023).

26. Miyazawa, K., et al. Cross-ancestry genome-wide analysis of atrial fibrillation unveils disease biology and enables cardioembolic risk prediction. Nat Genet 55, 187–197 (2023).

27. Lee, J.Y., et al. Korean atrial fibrillation network genome-wide association study for early-onset atrial fibrillation identifies novel susceptibility loci. Eur Heart J 38, 2586–2594 (2017).

28. Sakaue, S., et al. A cross-population atlas of genetic associations for 220 human phenotypes. Nat Genet 53, 1415–1424 (2021).

29. Depmap, B. DepMap 24Q2 Public. (10.25452/figshare.plus.25880521.v1, 2024).

30. Dempster, J.M., et al. Chronos: a cell population dynamics model of CRISPR experiments that improves inference of gene fitness effects. Genome Biol 22, 343 (2021).

31. Montgomery, P. Dependency Threshold. (DepMap Community Forum, https://forum.depmap.org/t/dependency-threshold/498, 2021).

32. Benjamini, Y. & Hochberg, Y. Controlling the False Discovery Rate: A Practical and Powerful Approach to Multiple Testing. Journal of the Royal Statistical Society Series B: Statistical Methodology 57, 289–300 (1995).

33. McFarland, J.M., et al. Improved estimation of cancer dependencies from large-scale RNAi screens using model-based normalization and data integration. Nat Commun 9, 4610 (2018).

34. Black, N., Mohammad, F., Saraf, K. & Morris, G. Endothelial function and atrial fibrillation: A missing piece of the puzzle? J Cardiovasc Electrophysiol 33, 109–116 (2022).

35. Tsunenari, T., et al. Structure-function analysis of the bestrophin family of anion channels. J Biol Chem 278, 41114–41125 (2003).

36. Yan, Z., Pu, X., Chang, X., Liu, Z. & Liu, R. Genetic basis and causal relationship between atrial fibrillation and sinus node dysfunction: Evidence from comprehensive genetic analysis. Int J Cardiol 418, 132609 (2024).

37. Song, W., Yang, Z. & He, B. Bestrophin 3 ameliorates TNFalpha-induced inflammation by inhibiting NF-kappaB activation in endothelial cells. PLoS One 9, e111093 (2014).

38. Deng, H., et al. Role of tumor necrosis factor-alpha in the pathogenesis of atrial fibrillation. Chin Med J (Engl) 124, 1976–1982 (2011).

39. Nso, N., Bookani, K.R., Metzl, M. & Radparvar, F. Role of inflammation in atrial fibrillation: A comprehensive review of current knowledge. J Arrhythm 37, 1–10 (2021).

40. Wang, X., et al. Genetic Susceptibility to Atrial Fibrillation Identified via Deep Learning of 12-Lead Electrocardiograms. Circ Genom Precis Med 16, 340–349 (2023).

41. Sau, A., et al. Artificial intelligence-enabled electrocardiogram for mortality and cardiovascular risk estimation: a model development and validation study. Lancet Digit Health 6, e791–e802 (2024).

42. Libiseller-Egger, J., et al. Deep learning-derived cardiovascular age shares a genetic basis with other cardiac phenotypes. Sci Rep 12, 22625 (2022).

43. Honda, M., et al. Vestigial-like 2 contributes to normal muscle fiber type distribution in mice. Sci Rep 7, 7168 (2017).

44. Song, K., et al. Heart repair by reprogramming non-myocytes with cardiac transcription factors. Nature 485, 599–604 (2012).

45. McFadden, D.G., et al. A GATA-dependent right ventricular enhancer controls dHAND transcription in the developing heart. Development 127, 5331–5341 (2000).

46. Morikawa, Y. & Cserjesi, P. Cardiac neural crest expression of Hand2 regulates outflow and second heart field development. Circ Res 103, 1422–1429 (2008).

47. Feghaly J, Zakka P, London B, MacRae Calum A, Refaat Marwan M. Genetics of Atrial Fibrillation. Journal of the American Heart Association. 2018;7(20):e009884.

48. Khawajakhail, R., et al. Advancements in gene therapy approaches for atrial fibrillation: targeted delivery, mechanistic insights and future prospects. Curr. Probl. Cardiol. 49, 102431 (2024).

